# Large-Scale Transcriptional Profiling of Molecular Perturbations Reveals Cell Type Specific Responses and Implications for Environmental Screening

**DOI:** 10.1101/2020.08.26.268458

**Authors:** Kun Zhang, Yanbin Zhao

## Abstract

Cell-based assays represent nearly half of all high-throughput screens currently conducted for risk assessment of environmental chemicals. However, the sensitivity and heterogeneity among cell lines has long been concerned but explored only in a limited manner. Here, we address this question by conducting a large scale transcriptomic analysis of the responses of discrete cell lines to specific small molecules. Our results illustrate heterogeneity of the extent and timing of responses among cell lines. Interestingly, high sensitivity and/or heterogeneity was found to be cell type-specific or universal depending on the different mechanism of actions of the compounds. Our data provide a novel insight into the understanding of cell-small molecule interactions and have substantial implications for the design, execution and interpretation of high-throughput screening assays.

## Introduction

Cell-based high-throughput screening (HTS) is extensively applied in environmental hazard and risk assessment. In the ToxCast program, approximately half of all >700 HTS assays are conducted in cells (1,2), and similar assays that act at the same gene or pathway targets are established in multiple cell types (2,3). For instance, human glucocorticoid receptor binding assays have been developed in A549, HepG2, MCF-7, U2OS, SH-SY5Y, PC3 and HEK293 cells (3–6). Consequently, sensitivity and heterogeneity of response among cell lines has long been a concern (5–8), but clear answers are still lacking due to limited comparative datasets.

Currently, the ongoing LINCS L1000 program (9), aims to collect molecular data from cells following exposure to thousands of molecules. This program provides a unique opportunity to uncover cell line specific responses. In our present analysis, we report on the collection of >60,500 gene expression signatures, composed of >223,300 gene expression profiles from a wide array of cell lines (derived from prostate, breast, lung, colon, liver, kidney, and skin cancer) exposed to 2243 molecules. This dataset covers >1300 pharmaceuticals currently on the market, of which over 900 have been detected in environmental waterbodies (see supporting data). The data bear information on fully annotated mechanisms of action in identical exposure procedures and each chemical was tested in at least seven of the nine most commonly employed HTS cell lines. With our analysis we aimed to address two principal questions: 1. Can cell-type specific responses and temporal variation in responses be observed at scale? 2. How can sensitivity and heterogeneity analyses among cell lines guide better environmental screening?

## Results and Discussion

Cell-type specific responses were represented by the coefficient of variation (CV) of the differentially expressed gene (DEGs) numbers and altered pathway numbers, which capture differences in both potency and efficacy (10). Standard deviation and mean value were provided for each of the 2243 molecules, as shown in Fig. 1A-B. Distinct responses among the nine cell lines tested were illustrated. More than 26% of signatures have CV>1 and 72% of signatures have CV>0.5 for the DEGs number values. For altered pathway numbers, they were at 81% and 96%, respectively. Temporal variations as assessed at 6-hour and 24-hour exposure were significantly different across all compounds exhibiting poor correlation with R2=0.039 (Fig. 1C), suggesting distinct transcriptional responses over time. Detailed pathway analyses demonstrated that 78% of signatures did not share any pathways (among the top 5) (Fig. 1D), further confirming the heterogeneity of response.

**Fig. 1.**
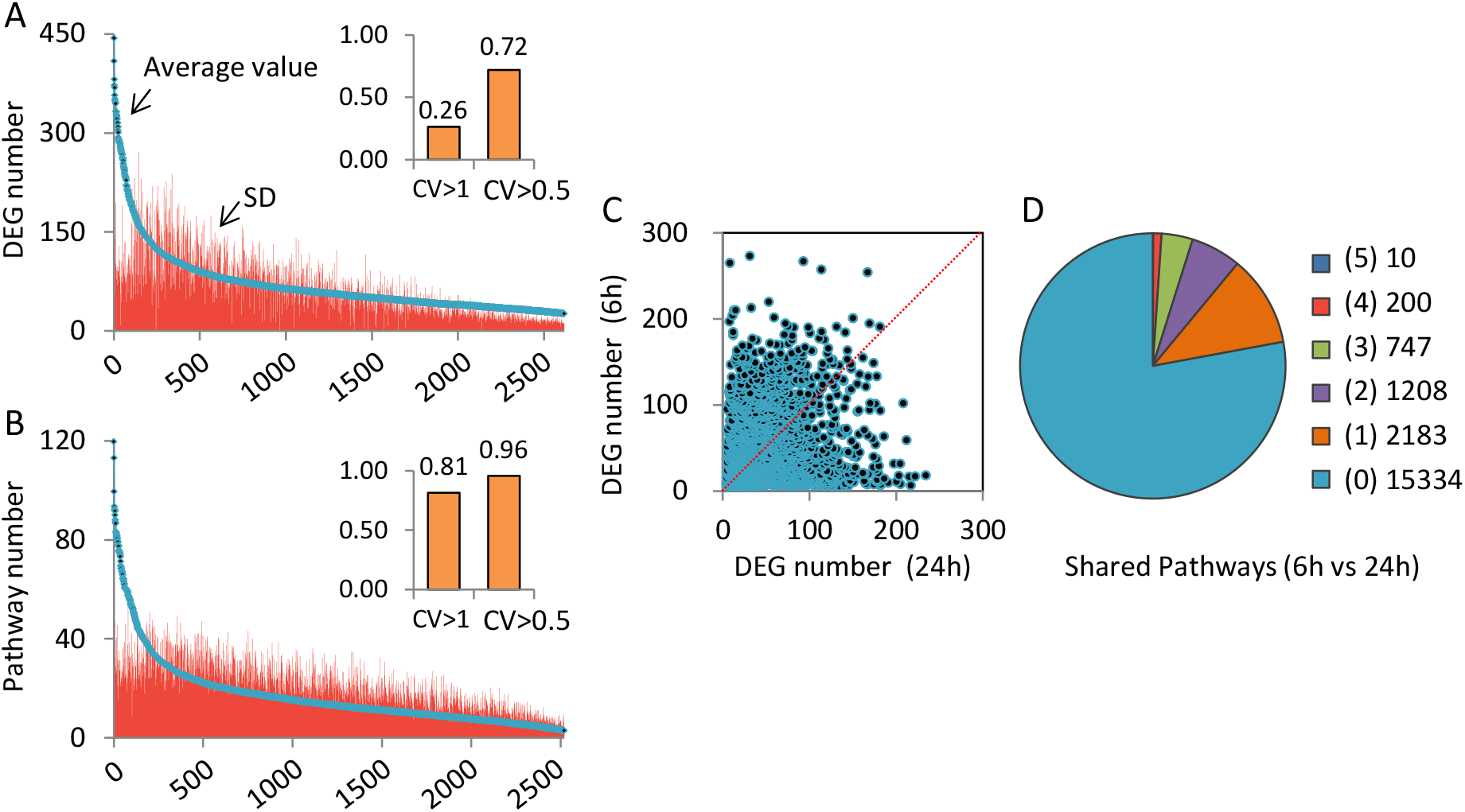
Cell-type specific responses and temporal variations. (A) The number of differentially expressed genes (DEGs) and standard deviation (SD) for each compound among nine cell lines analyzed. CV: coefficient of variation. (B) The number of altered pathways and standard deviation for each compound among nine cell lines analyzed. CV: coefficient of variation. (C) Correlations between the numbers of DEGs at 6-hour and 24-hour exposures. (D) Frequency of shared top 5 pathways between 6-hour and 24-hour exposures. Key on the right: numbers of compounds and numbers of shared pathways (inside bracket) among top 5 pathways generated via DAVID 6.8. For instance, (5)10 means that there were ten compounds for which the top 5 pathways were identical between 6-hour and 24-hour exposures.

Sensitivity and heterogeneity of response among different cell types to perturbations are of central importance in the performance of cell-based HTS. The SR ratio (defined as an exposed signature with greater DEG numbers than those in the related control groups, representing cell sensitivity) and E-index (representing cell heterogeneity) for each cell line and each of the twelve types of signal pathway modulators are provided in Fig. 2. In general, high sensitivity and/or heterogeneity are cell type-specific or universal depending on the mechanism of actions of the compounds. For glucocorticoid receptor agonists, high SR ratio (82% and 73%) and E-index (70% and 77%) are observed for HCC515 and A549, two lung carcinoma cells (Fig. 2A). The t-SNE plot further shows the distinct signatures clustering for them, which are remarkably different from those observed with other cell lines (Fig. 2B). For HDAC inhibitors, the very high SR ratio (ranged from 80% to 92%) and E-index (ranged from 82% to 94%) were universal among cell lines (Fig. 2A). The responses to HDAC inhibitors share very similar altered KEGG pathways (among the top 5), i.e. p53 signaling pathway and viral carcinogenesis (Fig. 2C). Similar results are found for ATPase inhibitors. Some pathway modulators, such as sodium channel blockers and calcium channel blockers, exhibit low SR ratios but high E-indices in some specific cell types (Fig. 2A). Figure 2D depicts the top two connections between cell line and each type of signal pathway modulators based on their SR ratio and E-index.

**Fig. 2.**
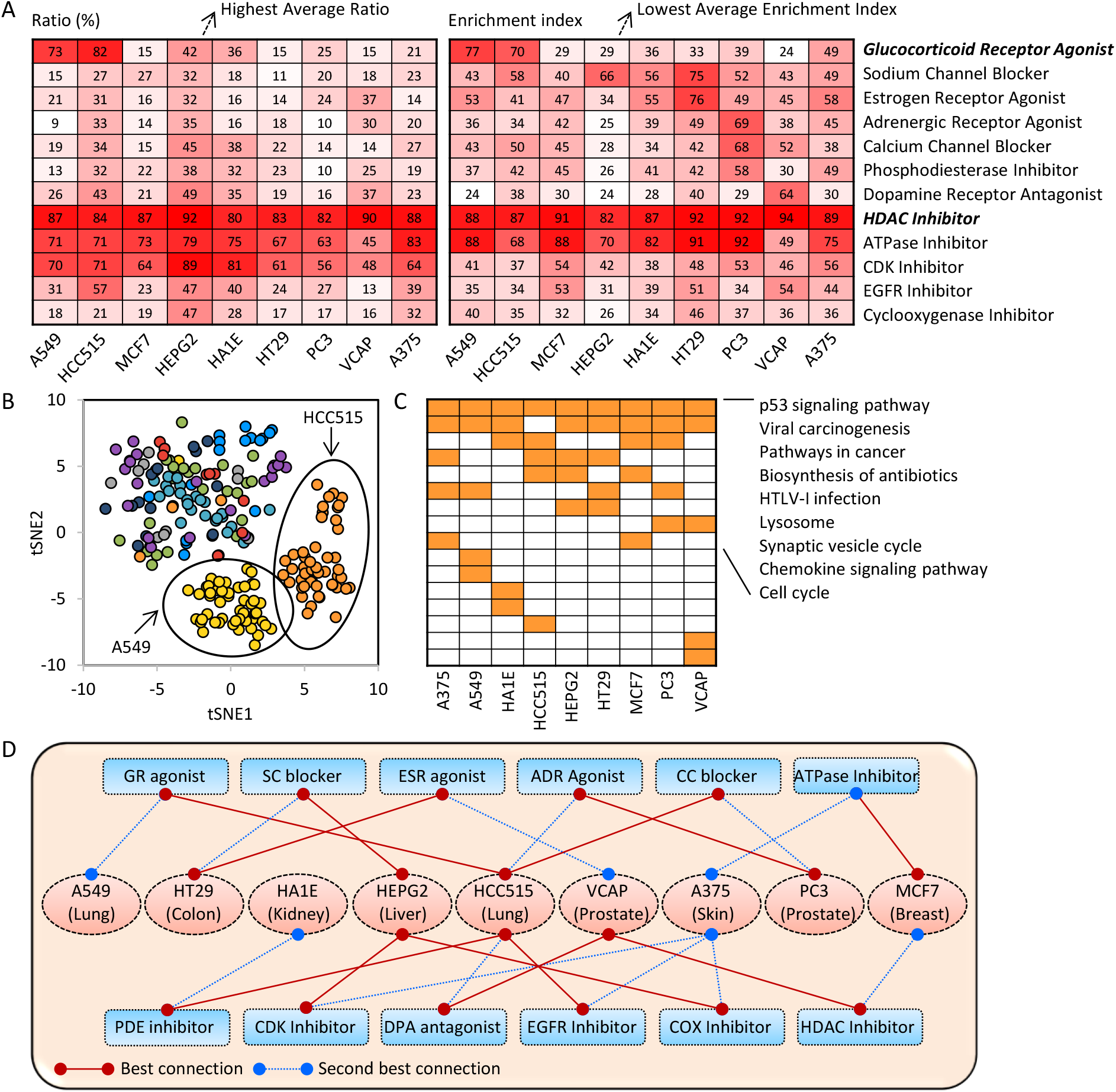
Sensitivity and heterogeneity among cell lines. (A) Scores of the significant response ratio and the enrichment index for each cell line and each of the twelve types of signal pathway modulators. The highest average ratio and lowest average enrichment index were observed for the HEPG2 cell line. (B) t-SNE plot showing the signature clustering for glucocorticoid receptor agonists. Colors represents different cell lines. (C) Shared top 5 pathways response to HDAC inhibitors exposure for each of the nine cell lines analyzed. Universal responses were observed for p53 signaling and viral carcinogenesis. (D) Schematic diagram depicts the best (solid red line) and second best (blue dotted line) connections between cell line and each type of signal pathway modulators.

Our analysis provides a novel and comprehensive perspective on cell-chemical interactions. It adds to a growing body of evidence suggesting that a large proportion of cell-type specific responses exist (8,11). We also observe remarkable temporal variations. Distinct transcriptional patterns occur for most chemicals at 6-hour compared with 24-hour exposures. Previous investigations have been inconsistent suggesting high consistency of gene expression over short time periods in some studies (12,13), while in others such consistency was not observed (14,15). Here, we demonstrate that for a very large proportion of chemicals, regardless of their mechanisms of action, little evidence of high temporal consistency can be found.

Despite the extent of cell type-specific and temporal variability, we attempted to identify the most suitable cells for HTS with a specific target. Answering this question cannot be straightforward, thus, we employed in the first instance twelve representative pathways extensively explored in environmental science (3). Our results reveal that for any given pathway, high heterogeneity and sensitivity to a chemical are readily classified as cell type-specific or universal. Lung carcinoma cells would be best suited for glucocorticoid receptor agonists. This is supported by previous binding sensitivity tests conducted in A549 and other cells (4–6). In contrast, for HDAC inhibitors and ATPase inhibitors, all cell lines show very high sensitivities and heterogeneities. HEPG2 cells have high sensitivity while low heterogeneity responses. This may be explained by complex metabolism process in hepatocytes, as described for some specific signal pathway modulators, such as RXR agonists (16).

Our results may have direct implications for the screening of environmental and other chemicals and raise questions regarding the design, execution and interpretation of HTS assays in the context of environmental risk assessment. For instance, is there a minimal number of cell types necessary to undertake a robust screen of small molecules? It would also be important to understand the extent to which any single cell line might be representative for screening environmental chemicals? Such questions will require a comprehensive exploration of chemical space using much larger libraries and much deeper pathway analyses.

## Materials and Methods

Raw gene expression profiles were obtained from the LINCS project generated by the Broad Institute (http://www.lincsproject.org/) (9). Characteristics of >19,800 related bioactive compounds, including their molecular features and mechanism of action (MOA) information, were obtained via the Drug Repurposing Hub (https://clue.io/repurposing) and the ChEMBL (https://www.ebi.ac.uk/chembl/). Four criteria to generate gene expression sub-profiles are proposed in the present study: fully annotated MOA information; testing performed in at least seven of the nine most commonly employed cell lines; time periods: 6 hour and 24 hour exposure; concentration: 10 μM. Finally, the data subset applied in the present study included 2243 molecules with 60,576 gene expression signatures that comprise >223,300 gene expression profiles.

The number of differentially expressed genes (DEGs) was computed via robust z-scores (9) for each of the 2243 molecules. Pathway enrichment analyses were batch processed via DAVID 6.8. Significant Response (SR) was defined as an exposed signature with greater DEG numbers than those in the related control groups. Afterwards, SR ratio (the number of significant response signatures/the number of total response signatures) was calculated for each of the nine cell lines and each of the twelve types of signal pathway modulators to represent cell line sensitivity. Enrichment index (E-index), representing the cell line heterogeneity, was estimated as:

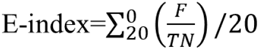

where F represents the most frequent DEGs shared among each type of significant response signatures, and TN represents the total number of exposure signatures for each cell line and each type of signal pathway modulators. The t-SNE plot was implemented using R (version 3.4.3) together with the Rtsne package. Additional information about methodology, research framework and supporting data were provided below.

## Author contributions

Y.B.Z. designed research; K.Z. performed research; Y.B.Z. wrote the paper.

## Notes

The authors declare no competing financial interest.

## Acknowledgments

We thank Calum MacRae (Brigham and Women’s Hospital, Harvard Medical School), Karl Fent (University of Applied Sciences and Arts Northwestern Switzerland and Swiss Federal Institute of Technology (ETH Zürich)) and Xuehan Zheng, Haochun Shi (Shanghai Jiao Tong University) for critically reading the manuscript and making valuable suggestions. This work was funded by the Shanghai Pujiang Program [19PJ1404800] and the Startup Fund for Youngman Research at Shanghai Jiao Tong University [WF220416007 and WF112116003].

## Supplementary Information

We provide the following supplementary information that the editors might consider useful for the review. If published, this information will be available on the authors’ own website.

Details of datasets and research framework information (Figure S1); classification of target 2243 compounds (Figure S2); STP (shared top pathways) index (Figure S3); differentially expressed genes and pathways for dexamethasone (DEX) and betamethasone (BET) (Figure S4); differentially expressed genes and pathways for nifedipine (NIF) and nimodipine (NIM) (Figure S5); 401 compounds information for 12 target pathways analyzed (Table S1); shared differentially expressed genes and pathways between DEX and BET for HCC515 and A549 cell lines (Figure S6); average ratio and enrichment index for the nine target cell lines (Figure S7); characteristics for sodium channel blocker and calcium channel blocker (Figure S8).

**Figure S1.**
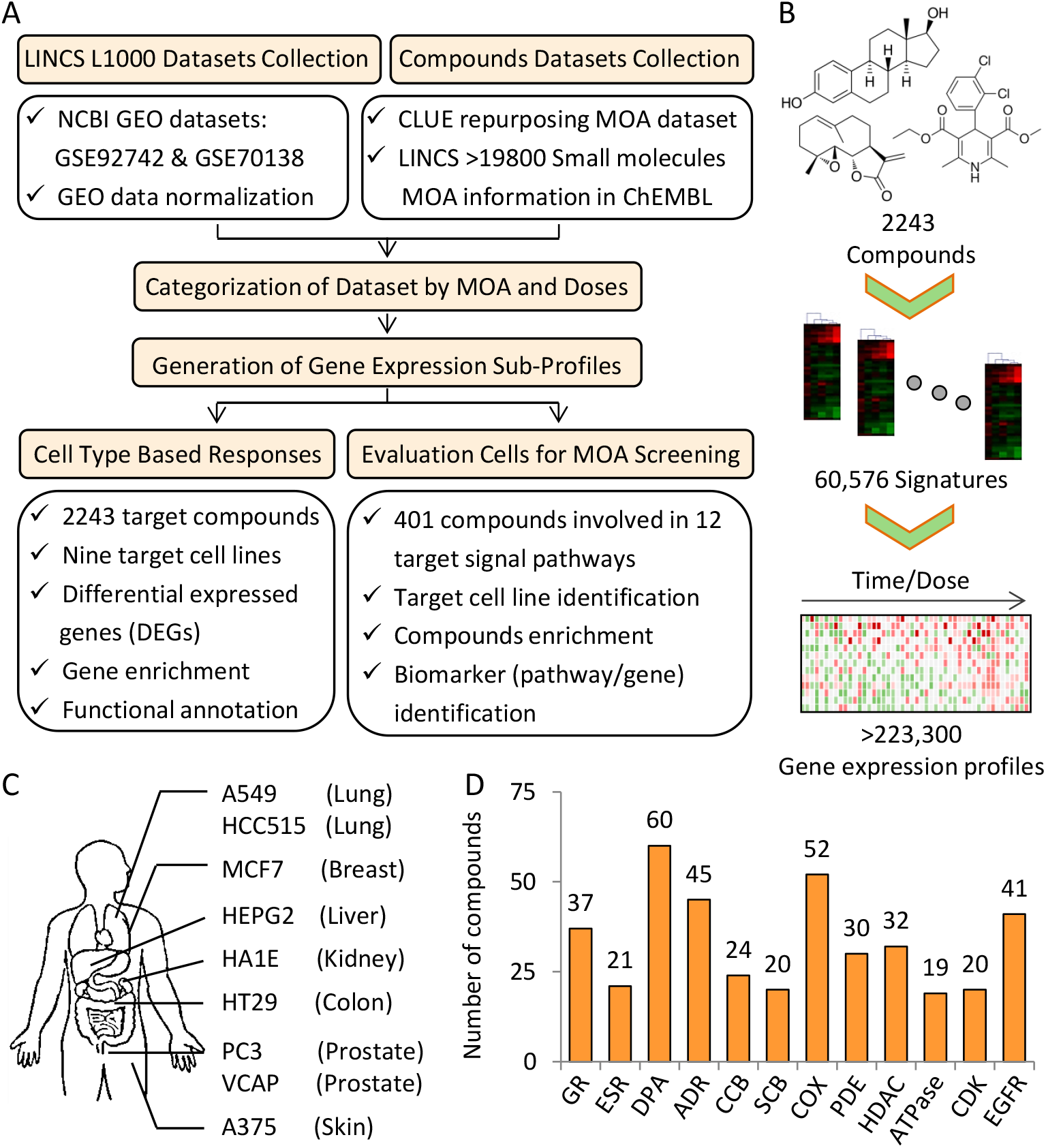
Datasets and research framework of the present analysis. (A) Datasets collection and detailed workflow. (B) Sub-datasets employed in the present study. (C) Nine cell lines including A549, HCC515, MCF7, HEPG2, HA1E, HT29, PC3, VCAP and A375 analyzed in the present study. (D) Twelve representative pathways, covered 401 modulators, that were extensively explored for sensitivity and heterogeneity analysis.

**Figure S2.**
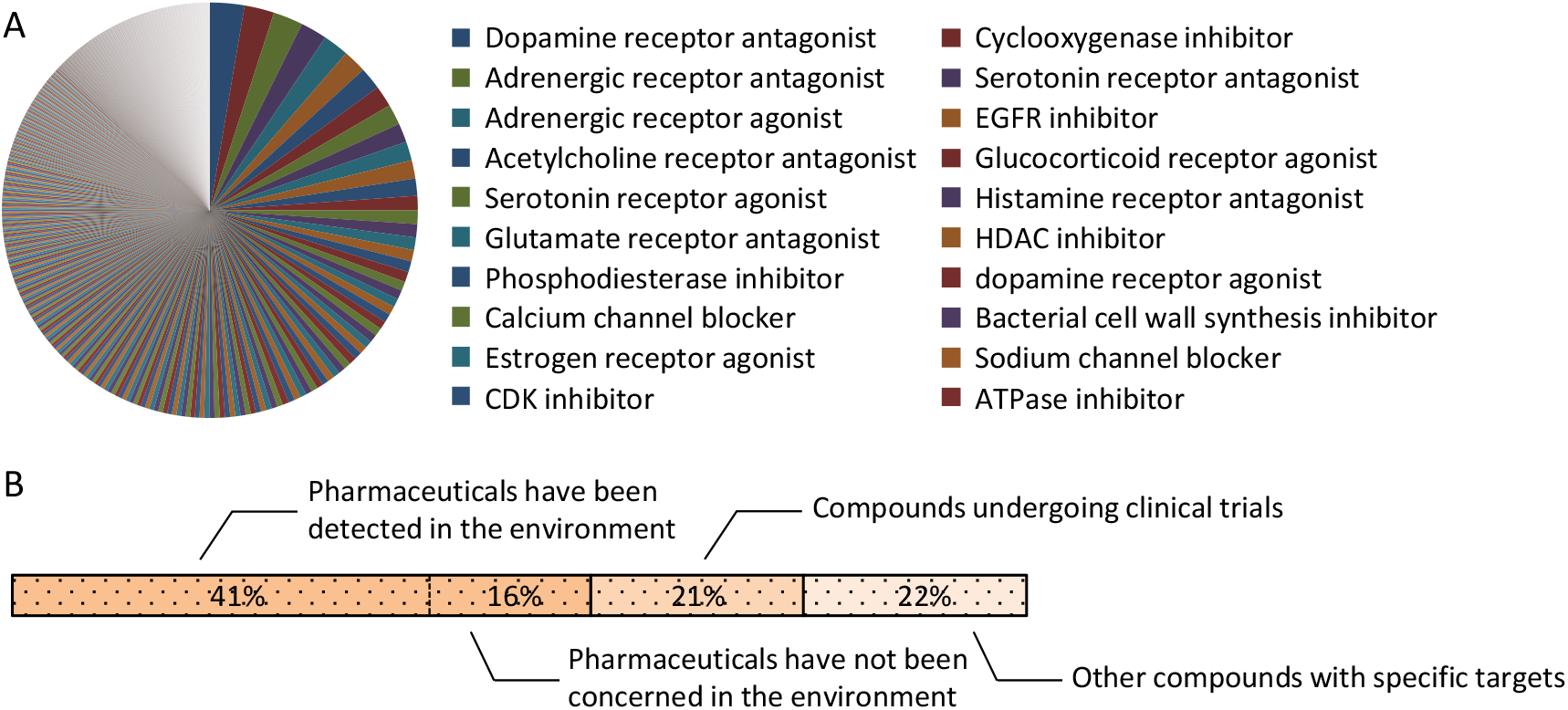
Classification of the target 2243 compounds analyzed in the present study. (A) Clustering showing different mechanism of actions. (B) Clustering showing environmental occurrence.

**Figure S3.**
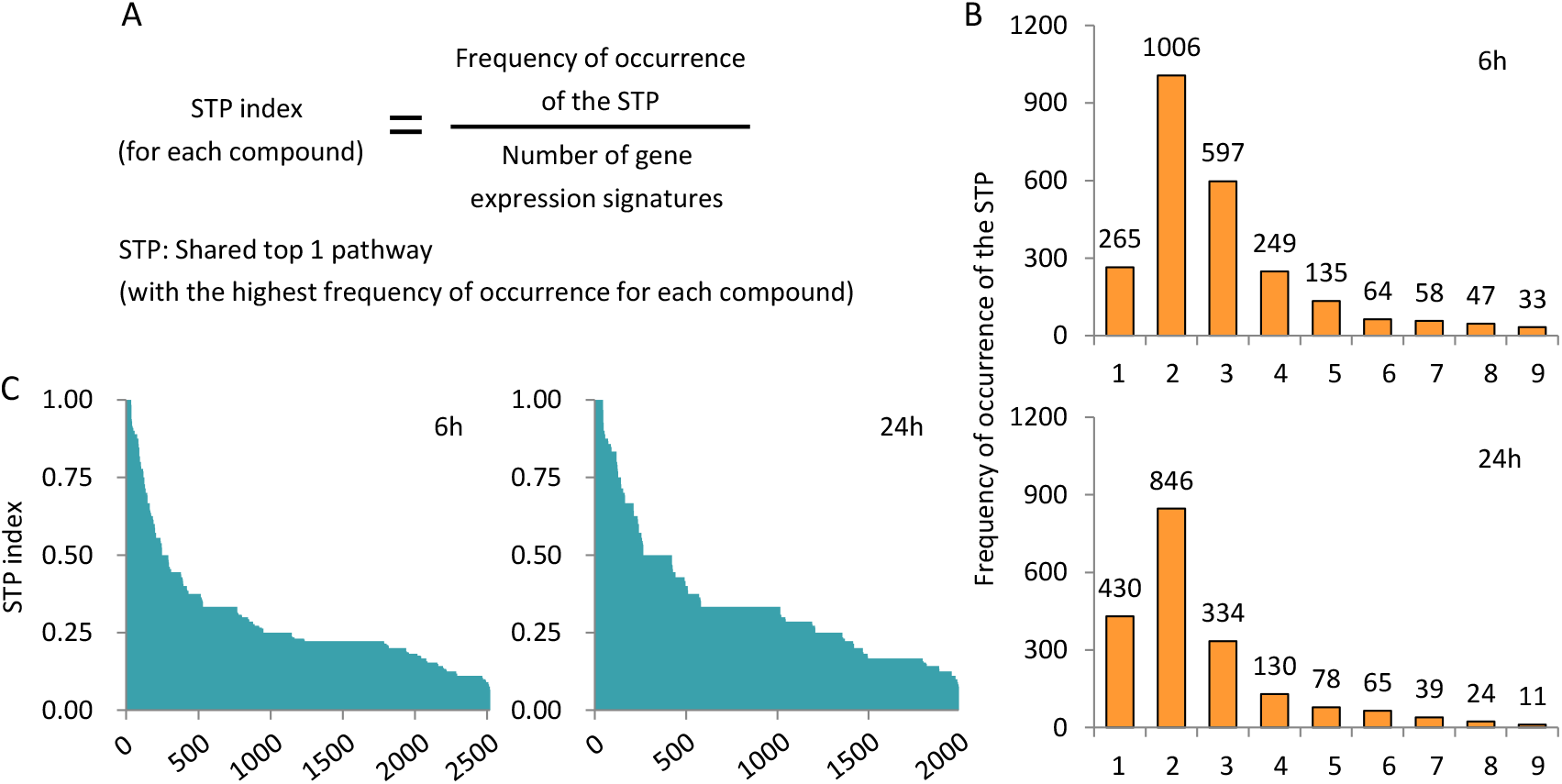
STP (shared top 1 pathway) index for each compound. (A) Definition of the STP index. (B) The frequency of occurrence of STP among nine cell lines analyzed. (C) Average STP index for each of the target 2243 compounds analyzed.

**Figure S4.**
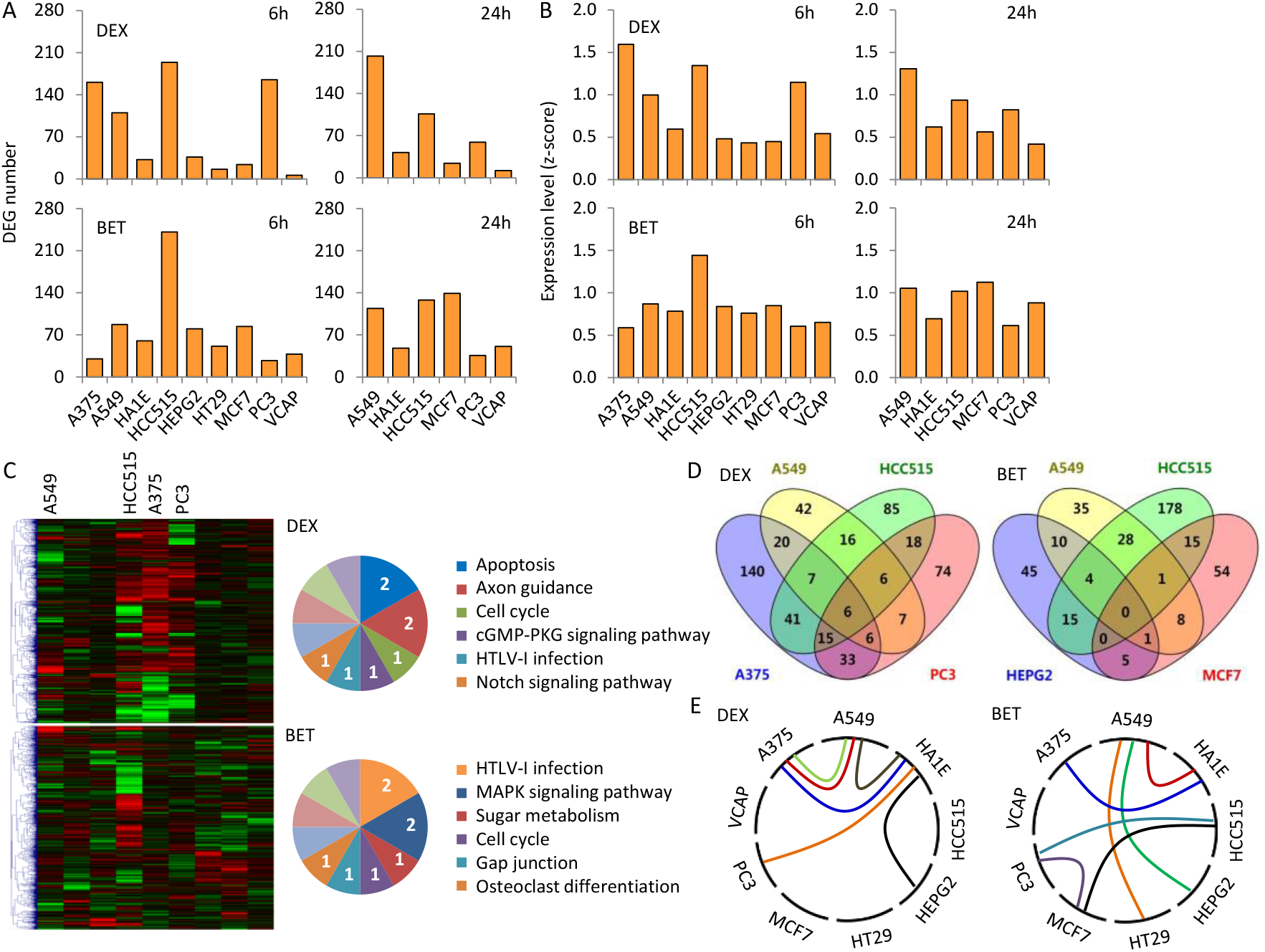
Sensitivity and heterogeneity analysis among cells for glucocorticoid receptor agonists, dexamethasone (DEX) and betamethasone (BET). (A) Differences in the numbers of differentially expressed genes (DEGs). (B) Differences in the average expression levels. (C) Clustering map represents the differences in pathways. (D) Limited shared DEGs among cell lines. (E) Limited shared pathways (among top 5) among cell lines.

**Figure S5.**
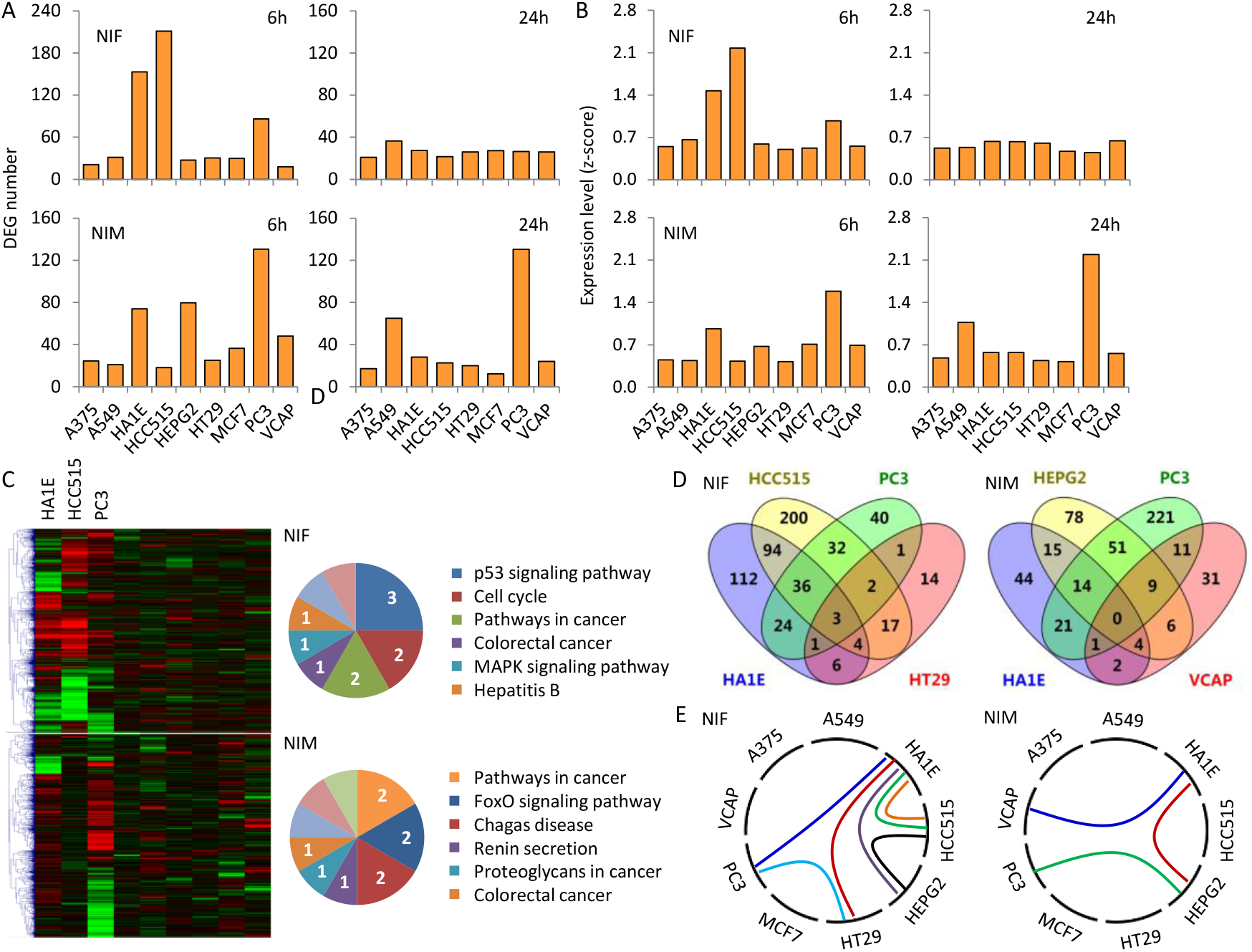
Sensitivity and heterogeneity analysis among cells for calcium channel blockers, nifedipine (NIF) and nimodipine (NIM). (A) Differences in the numbers of differentially expressed genes (DEGs). (B) Differences in the average expression levels. (C) Clustering map represents the differences in pathways. (D) Limited shared DEGs among cell lines. (E) Limited shared pathways (among top 5) among cell lines.

**Table S1.**
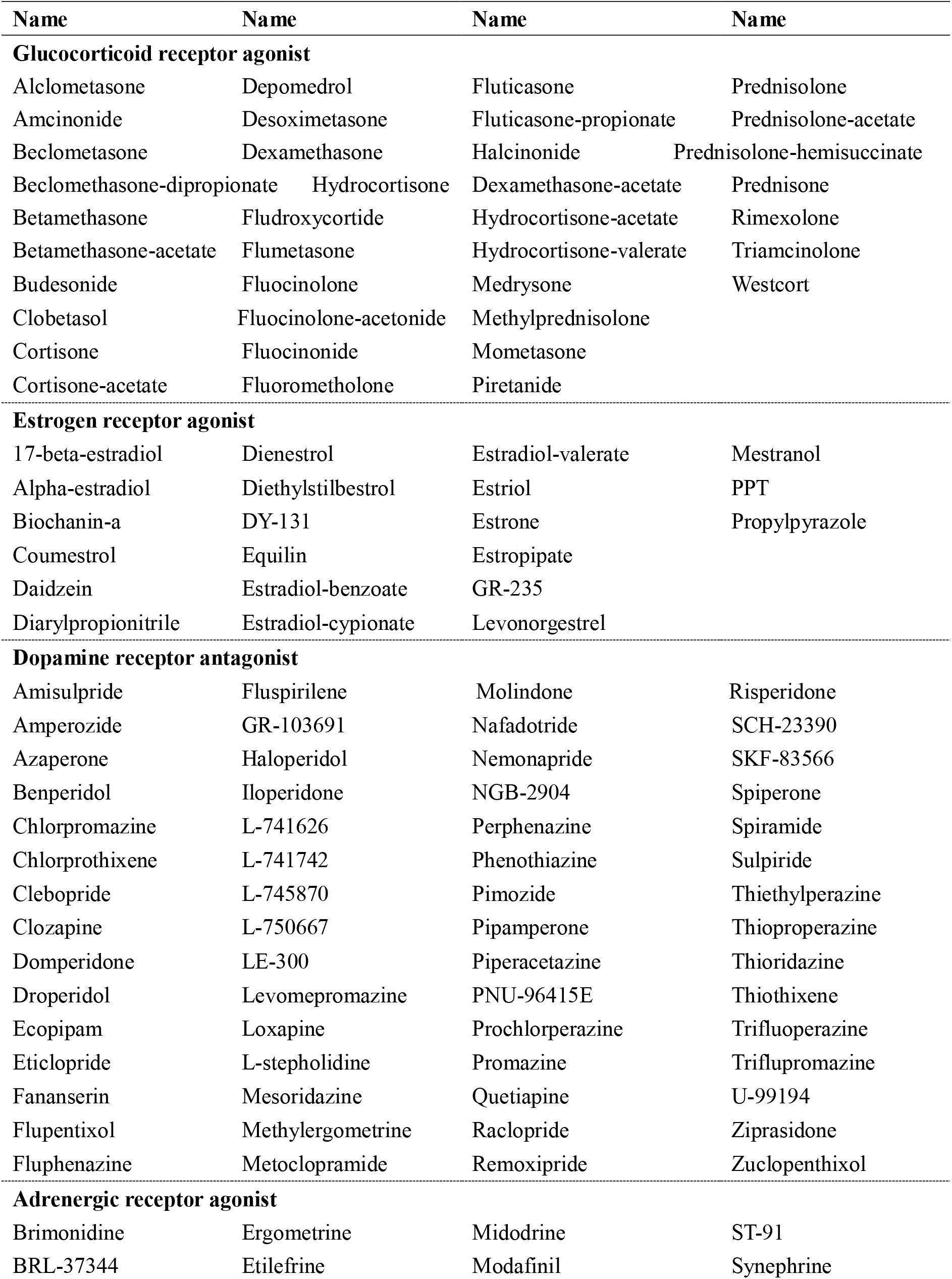
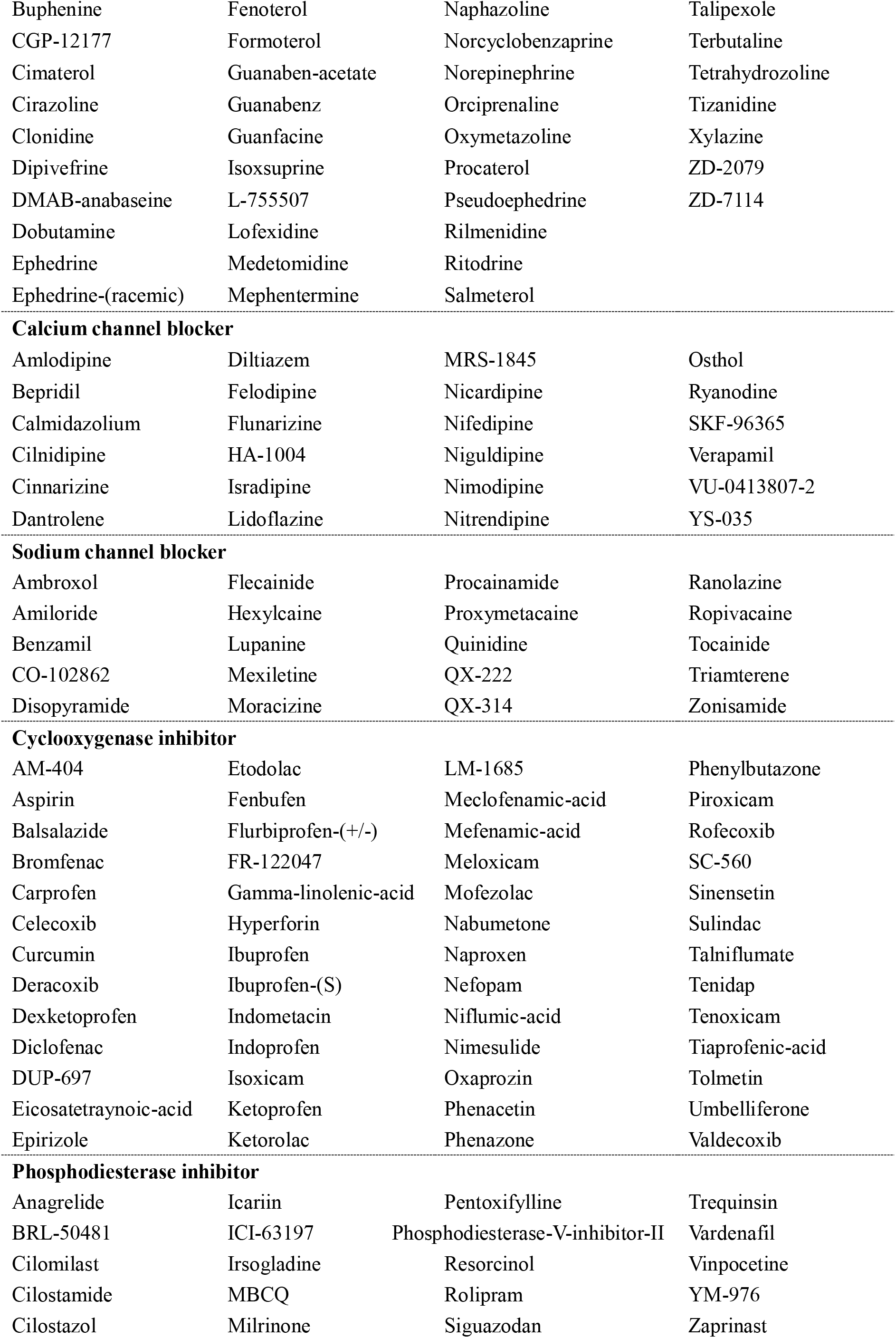
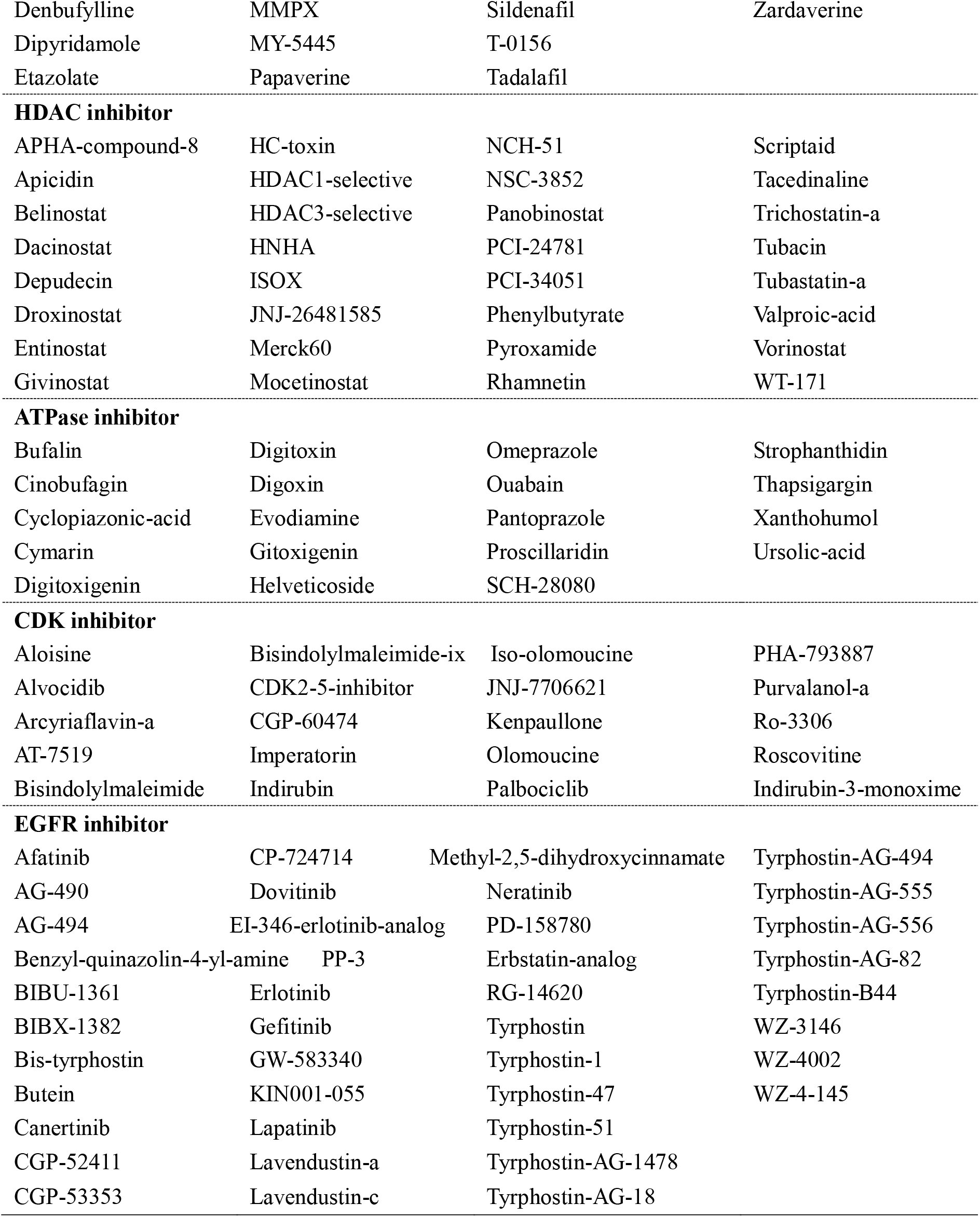
Classification of the 401 compounds involved in 12 target pathways analyzed in the present study. They included: glucocorticoid receptor agonist; estrogen receptor agonist; dopamine receptor antagonist; adrenergic receptor agonist; calcium channel blocker; sodium channel blocker; cyclooxygenase inhibitor; phosphodiesterase inhibitor; HDAC inhibitor; ATPase inhibitor; CDK inhibitor and EGFR inhibitor.

**Figure S6.**
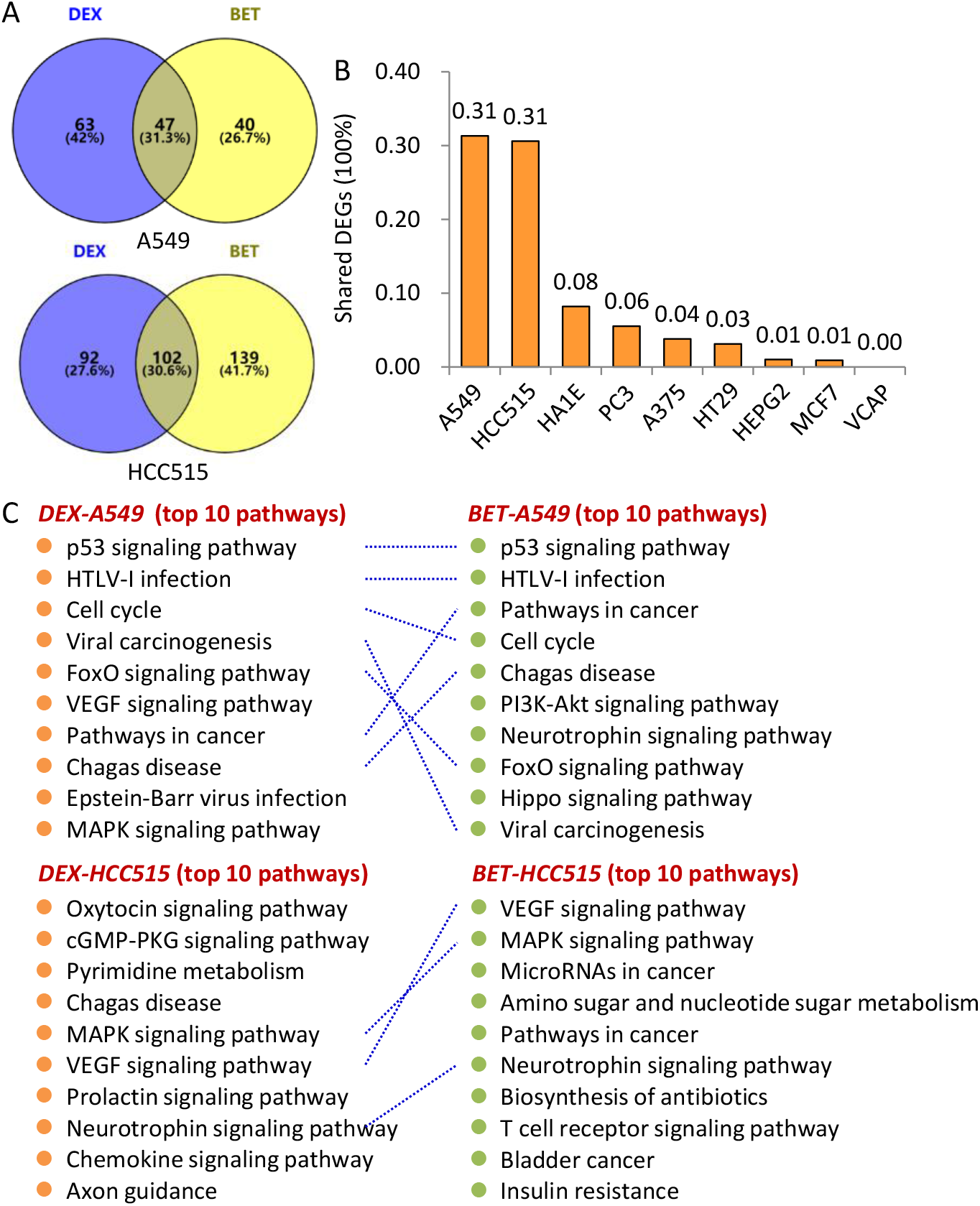
Similarity analysis between DEX and BET. (A-B) Shared differentially expressed genes between DEX and BET for HCC515 and A549 cell lines. (C) Shared altered pathways between DEX and BET for HCC515 and A549 cell lines.

**Figure S7.**
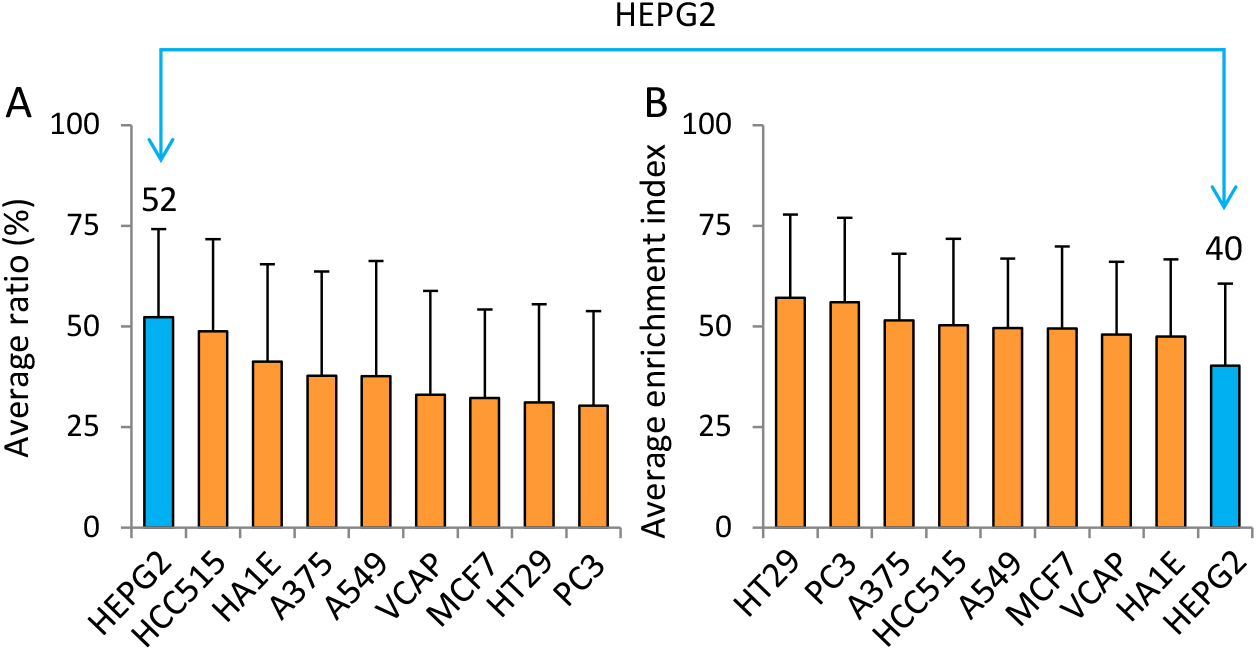
Average significant response ratio (SR ratio) and enrichment index (E-index) for each of the nine cell lines analyzed. The highest average SR ratio and the lowest average E-index were observed for HEPG2 cells.

**Figure S8.**
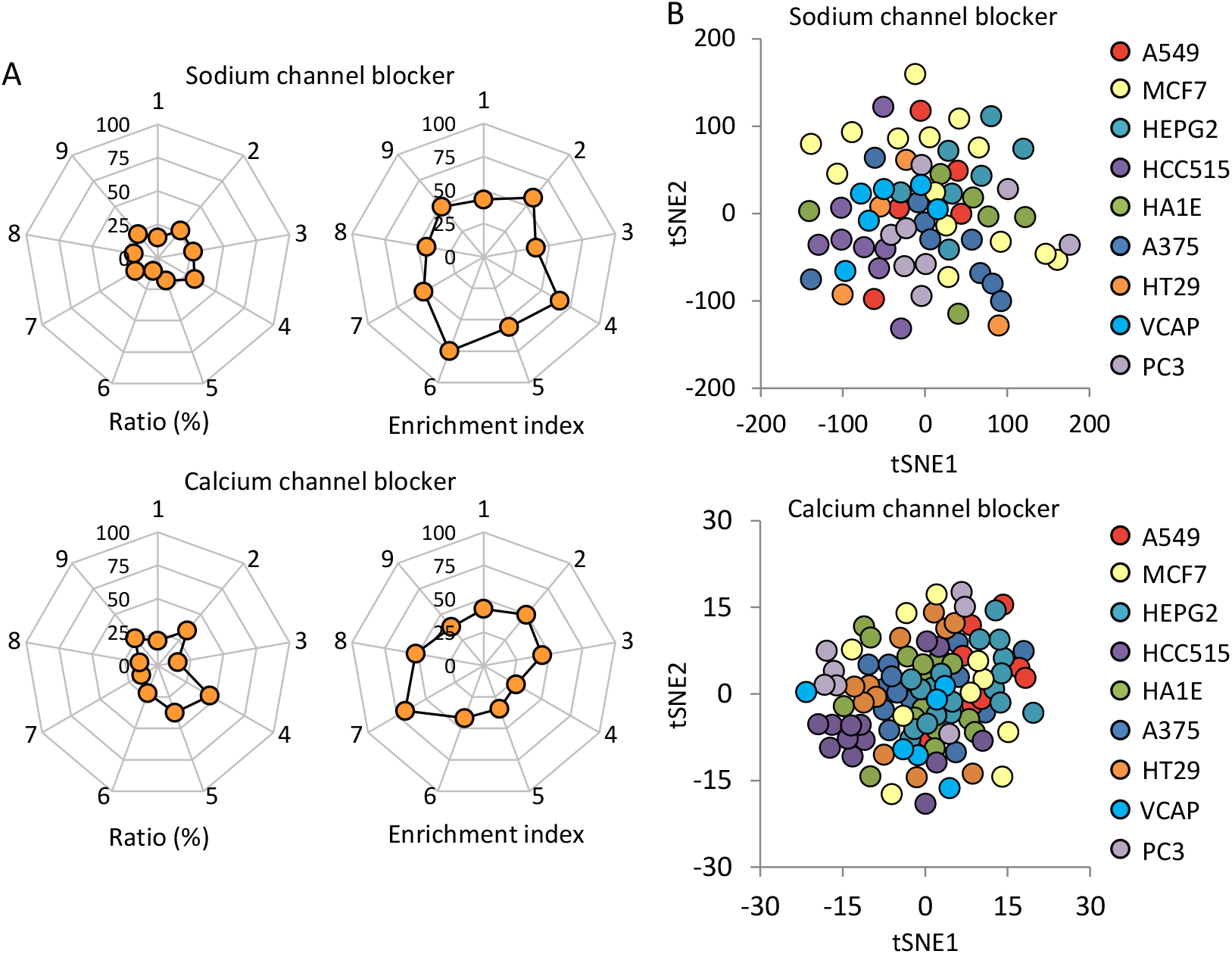
Characteristics for sodium channel blockers and calcium channel blockers. (A) SR ratio and E-index for each of the nine cell lines analyzed. Key: 1-9 represent the A549, HCC515, MCF7, HEPG2, HA1E, HT29, PC3, VCAP and A375 cell lines. (B) t-SNE plots visualizing cluster assignments of cells. Colors represents the different cell lines.

## References

(1) R. Tice, C. Austin, R. Kavlock, J. Bucher, Improving the human hazard characterization of chemicals: a Tox21 update. Environ. Health Perspect. 121, 756–765 (2013).

(2) R. S. Judson, et al., In vitro screening of environmental chemicals for targeted testing prioritization: the ToxCast project. Environ. Health Perspect. 118, 485–492 (2010).

(3) S. Kim, et al., PubChem 2019 update: improved access to chemical data. Nucleic. Acids Res. 47, D1102–D1109 (2019).

(4) M. Schriks, et al., High-resolution mass spectrometric identification and quantification of glucocorticoid compounds in various wastewaters in the Netherlands. Environ. Sci. Technol. 44, 4766–4774 (2010).

(5) H. W. Xiao, et al., Haploinsufficiency of NR3C1 drives glucocorticoid resistance in adult acute lymphoblastic leukemia cells by down-regulating the mitochondrial apoptosis axis, and is sensitive to Bcl-2 blockage. Cancer Cell Int. 19, 218 (2019).

(6) W. W. Lockwood, K. Zejnullahu, J. E. Bradner, H. Varmus, Sensitivity of human lung adenocarcinoma cell lines to targeted inhibition of BET epigenetic signaling proteins. Proc. Natl. Acad. Sci. U S A. 109, 19408–19413 (2012).

(7) S. Jaeger, M. D. Frigola, P. Aloy, et al., Drug sensitivity in cancer cell lines is not tissue-specific. Mol. Cancer. 14, 40 (2015).

(8) J. Barretina, et al., The Cancer Cell Line Encyclopedia enables predictive modelling of anticancer drug sensitivity. Nature. 483, 603–607 (2012).

(9) A. Subramanian et al., A next generation connectivity map: L1000 platform and the first 1,000,000 profiles. Cell. 171, 1437–1452 (2017).

(10) M. Niepel, et al., Common and cell-type specific responses to anti-cancer drugs revealed by high throughput transcript profiling. Nat. Commun. 8 (1), 1186 (2017).

(11) M. A. Troester, et al., Cell-type-specific responses to chemotherapeutics in breast cancer. Cancer Res. 64, 4218–4226 (2004).

(12) L. M. Beaver, et al., Transcriptome analysis reveals a dynamic and differential transcriptional response to sulforaphane in normal and prostate cancer cells and suggests a role for Sp1 in chemoprevention. Mol. Nutr. Food Res. 58, 2001–2013 (2014).

(13) M. L. Kwok, X. L. Hu, Q. Meng, K. M. Chan, Whole-transcriptome sequencing (RNA-seq) analyses of the zebrafish liver cell line, ZFL, after acute exposure to Cu 2+ ions. Metallomics. 12, 732–751 (2020).

(14) Y. S. Zhang, et al., Estrogen induces dynamic ERα and RING1B recruitment to control gene and enhancer activities in luminal breast cancer. Sci. Adv. 6, eaaz7249 (2020).

(15) H. Ma, et al., Transcriptome analysis of glioma cells for the dynamic response to γ-irradiation and dual regulation of apoptosis genes: a new insight into radiotherapy for glioblastomas. Cell Death Dis. 4, e895 (2013).

(16) P. T. Vedell, et al., Global molecular changes in rat lives treated with RXR agonists: a comparison using transcriptomics and proteomics. Pharma. Res. Per. 2, e00074 (2014).

